# metagRoot: A comprehensive database of protein families associated with plant root microbiomes

**DOI:** 10.1101/2025.05.22.653656

**Authors:** Maria N. Chasapi, Iro N. Chasapi, Eleni Aplakidou, Fotis A. Baltoumas, Evangelos Karatzas, Ioannis Iliopoulos, Dimitrios J. Stravopodis, Ioannis Z. Emiris, Aydin Buluç, Ilias Georgakopoulos-Soares, Nikos C. Kyrpides, Georgios A. Pavlopoulos

## Abstract

The plant root microbiome is vital in plant health, nutrient uptake, and environmental resilience. To explore and harness this diversity, we present metagRoot, a specialized and enriched database focused on the protein families of the plant root microbiome. MetagRoot integrates metagenomic, metatranscriptomic, and reference genome-derived protein data to characterize 71,091 enriched protein families, each containing at least 100 sequences. These families are annotated with multiple sequence alignments, CRISPR elements, Hidden Markov Models, taxonomic and functional classifications, ecosystem and geolocation metadata, and predicted 3D structures using AlphaFold2. MetagRoot is a powerful tool for decoding the molecular landscape of root-associated microbial communities and advancing microbiome-informed agricultural practices by enriching protein family information with ecological and structural context. The database is available at https://pavlopoulos-lab.org/metagroot/ or https://www.metagroot.org

## Introduction

The discovery of the roles of proteins whose expression takes place in distinct plant microenvironments, such as aerial roots, bulbs, endosphere, nodules, rhizoplane, rhizosphere, stems, and stem tubers is crucial for the continuous progress of plant biology. These compartments are the hosts of dissimilar groups of proteins associated with nutrient transport, signal responses, plant-microbe stress reactions, and vegetation-plant interactions. Instead of being evenly distributed throughout the plant, most of these proteins are spatially restricted, pointing to each environment subtype’s unique physiological and ecological roles. For example, proteins involved in nitrogen metabolism are primarily located in nodules. However, those associated with reactive oxygen species signaling, redox regulation, and cell wall remodeling are mainly found in rhizoplane and rhizosphere, where a plant is exposed to soil microbes and abiotic stressors (1).

Plant compartments provide a biological habitat for diverse microorganisms (bacteria, fungi, archaea) closely linked with plant proteins to govern growth, immunity, and metabolism. For example, the secretion of host proteins, including expansins, defensins, and peroxidases, can alter the microbial composition and activity in the endospheres and rhizoplane. As a result, these plant proteins’ expression and activity can be changed by microbial metabolites, generating a well-regulated system and a great deal of dynamics. These relationships are not merely beneficial but often essential. Symbiotic microbes in nodules, for example, provide fixed nitrogen, while root-associated microbiota can enhance phosphate solubilization, stress tolerance, and pathogen resistance through coordinated interactions with plant proteins (2–5).

Generic protein and protein family databases such as UniProtKB (6), Protein Data Bank (PDB) (7), RefSeq (8), GenBank (9), IMG/M (10), Uniparc (6), Big Fantastic Database (BFD) (11), MGnify (12), Pfam(13), InterPro(14), SMART(15), NMPFamsDB (16, 17), COGs (18) and others (19–21) provide curated information on protein function, structural information, and interactions for most reference and metagenomic proteomes. Meanwhile, emerging resources like CyanoBase and RhizoBase (22), as well as the Microbial Genomes Atlas (MiGA) (23), facilitate cross-referencing between plant proteomic data and the taxonomic and functional profiles of associated microbes. However, to our knowledge, no existing database integrates reference proteomes with metagenomic and metatranscriptomic proteins at the protein family level for plant roots and their subhabitats. By examining proteins and protein families unique to specific plant environments or shared across certain subhabitats, researchers can delve deeper into the functional diversity underpinning plant adaptation and health, potentially uncovering new molecular functions and biological processes. In light of this, we introduce metagRoot, a comprehensive catalog of microbial protein families specific to plant root compartments.

## Materials and methods

### Data collection, filtering, and protein family generation

In October 2023, all publicly accessible plant root-associated metagenomes, metatranscriptomes, and reference genomes were sourced from the Integrated Microbial Genomes & Microbiomes (IMG/M) database (10), maintained by the Joint Genome Institute. The compiled dataset included 1,280 assembled metagenomes and 335 metatranscriptomes, collectively containing 3,497,616,101 protein-coding sequences and 3,207,863,396 scaffolds. Filtering criteria were applied to ensure a dataset of high-quality protein predictions (Figure 1) (17). To reduce truncation risk, sequences within 10 nucleotides of scaffold ends were removed. Additionally, only scaffolds longer than 500 nucleotides were retained to ensure that mainly complete genes were represented. Low-complexity regions were masked using the tantan application (version 26) to further refine the dataset. Any remaining sequences shorter than 35 amino acids were discarded. From this dataset, only samples linked to aerial roots, nodules, rhizoplane, endosphere, rhizosphere, stems, and stem tubers according to GOLD classification ecosystem subtypes (24), comprising 1,527 datasets (1,199 metagenomes, 327 metatranscriptomes), with 29,019,966 scaffolds and 403,495,896 proteins. For reference genomes, 17,825,681 protein sequences were extracted from 2,978 bacterial isolate genomes and 2,428 sequences from one root-associated archaeal isolate genome, yielding 2,979 datasets and a total of 421,324,005 sequences in the final catalog. For protein family generation, the MMseqs2 (25) Linclust (26) clustering algorithm was executed in bidirectional mode with parameters set to 30% sequence identity and 80% coverage. Protein families with 100 or more members were selected and aligned using MAFFT (27). During alignment, trimming was applied to minimize gaps. After aligning the sequences, the hhfilter (28) was employed to reduce redundancy within each family by applying a 95% sequence identity threshold and a 70% coverage criterion. This step minimizes bias from highly similar sequences and improves the accuracy of subsequent structural predictions (see below). To preserve biodiversity and ensure rich informational content, only families that originally contained 100 or more members before applying the hhfilter were reported. The final dataset consists of 71,091 families from 4,480 datasets, 25,961,294 (metagenome) scaffolds, 3,070,323 reference genomes, and 38,700,093 proteins.

**Figure 1.**
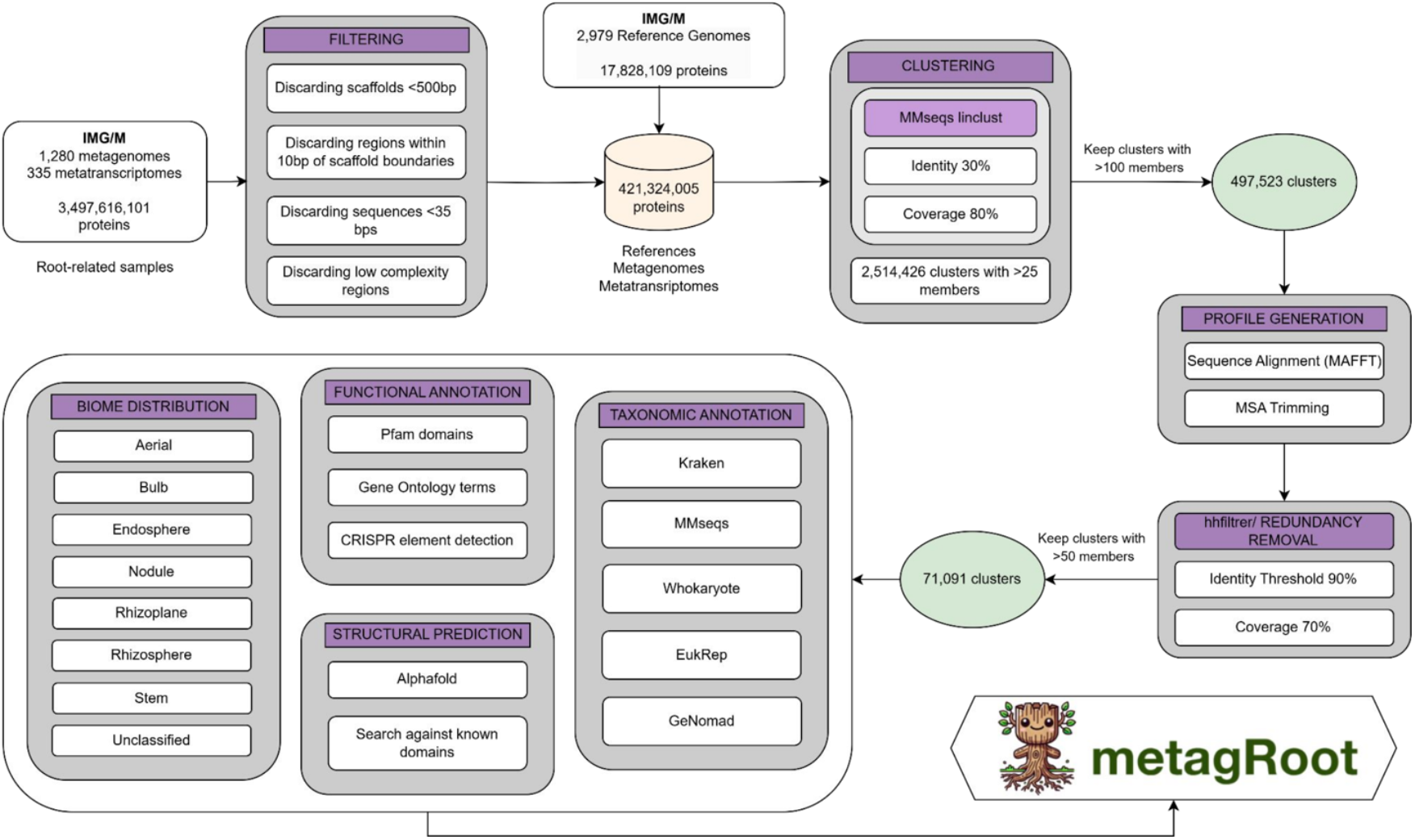
Data preprocessing and filtering. Pipeline illustrating the sequence filtering and clustering methodology for generating protein families from both reference genomes and metagenomes/metatranscriptomes.

### Taxonomy

The taxonomic analysis focused on protein families with more than 100 members to ensure statistical robustness. The scaffold sequences corresponding to protein families were retrieved from the IMG/M database. Classification began with Kraken2 (29), known for its high accuracy, and was further refined by applying MMseqs taxonomy (30) to annotate eukaryotic proteins better and reduce false positives. For scaffolds that remained unclassified, Whokaryote (31) was used to differentiate eukaryotic from prokaryotic contigs based on domain-specific gene structural features, followed by EukRep (32) to detect eukaryotic sequences further. Additionally, GeNomad (33) was employed to identify plasmid and viral sequences, ensuring a comprehensive assignment of mobile genetic elements. Ultimately, this multi-tool approach resulted in 37,155,632 scaffolds classified as Bacteria, 63,417 as Archaea, 45,286 as Eukaryota, 15,844 as Viruses, 6,582 identified as plasmid-derived, two linked to organelles, one classified as synthetic, and 1,387,727 remaining unclassified.

### Biome and metadata collection

Insights into habitat-specific diversity were obtained by examining the distribution of these protein families across various ecosystems, including aerial roots, bulbs, nodules, the rhizoplane, endosphere, rhizosphere, stems, and stem tubers, as defined by the GOLD classification. Environmental metadata were extracted from the IMG/M database and organized using the GOLD taxonomy, which categorizes ecosystems hierarchically into Environmental, Engineered, and Host-Associated groups. Within the Host-Associated microbiome category, a focus was placed on root-associated ecosystem subtypes. Within the Host-Associated microbiome category, root-associated ecosystem subtypes were prioritized, with dataset distribution prior to taxonomic family classification showing 1,898 rhizosphere entries, followed by 1,209 unclassified root-associated systems, 970 nodule samples, 199 rhizoplane datasets, 188 endosphere records, 29 aerial root collections, 10 stem tuber examples, and one dataset each documenting bulbs and stems. After family generation, among the 71,091 families containing 100 or more members, the composition was slightly adjusted to 1,879 rhizosphere datasets, 1,205 unclassified root-associated datasets, 970 nodule datasets, 199 rhizoplane datasets, and 186 endosphere datasets. Additionally, fewer datasets remained from other ecosystem subtypes, including 29 aerial root samples, 10 stem tuber samples, and one sample each from bulbs and stems.

### Functional annotation and CRISPR elements

Protein families were annotated by searching the representative sequences against Pfam (v37, Dec 2024), which includes 21,979 families and 709 clans, using hmmsearch (HMMER 3.3.2) with the trusted cutoff. In total, 66,854 families were successfully annotated with Pfam hits. CRISPR-related families were identified by first detecting CRISPR arrays using CRT (CRISPR Recognition Tool) (34), which recognizes direct repeats and spacers. In the total dataset of 25,961,294 scaffolds from metagenomes and metatranscriptomes in the 71,091 families, 8,927 were found with CRISPR arrays. Additionally, the protein families were screened for CRISPR-associated (Cas) proteins and other functional components based on Pfam hits. To this end, 1,223 scaffolds were detected with CRISPR Cas elements. At a family level, 24,078 of them were found to contain CRISPR arrays, whereas 6 families were annotated for Cas systems based on the CRISPR-associated Pfam domains. Collectively, 6 families were found to have a complete CAS system (both arrays and Cas elements).

### Structural model prediction

Structural information serves as a critical complement to sequence analysis, either confirming predicted functional elements by revealing their three-dimensional organization or refining annotations that sequence data alone cannot resolve. To this end, AlphaFold2 was used to predict structures for protein families with at least 50 effective sequences (upon hhfiltering for redundancy removal. For each family, five models were generated, and the one with the highest pLDDT score was selected. Only models classified as high (pTM ≥0.7) or medium-quality (pTM between 0.5 and 0.7) were retained. These models were then compared to experimentally validated structures from CATH (v4.4, Feb 2023) and PDB (2024 release), as well as both experimental and predicted structures in AlphaFoldDB, using Foldseek (35). Matches with a TM-score greater than 0.5 were considered valid. For families that lacked TM-alignment hits, additional criteria were applied to ensure alignment relevance. If the query was shorter than the target, the query alignment score had to be above 0.5. Similarly, if the query was longer, at least 50% of the target had to be aligned. Finally, superfamilies were defined using Foldseek in bidirectional mode, grouping structures based on 80% coverage and a TM-score of at least 0.5. The final dataset consisted of 71,088 family structural models, with 59,059 classified as High Quality and 9,036 as Medium Quality, and subsequently grouped into 9,463 superfamilies. Comparative results and structural analyses with CATH (36), PDB (7), and AlphaFold (37) revealed a total of 66,837 hits in CATH, including four unique entries; 66,910 hits in PDB, with no unique entries; and 68,049 hits in AlphaFold, with 624 unique entries. The analysis led to the discovery of 28 novel structures.

### Database Implementation and Structure

The metagRoot database uses MySQL as its main database management system and is built on the ASP.NET Core MVC framework. The back-end is coded in C#, while the front-end employs HTML, CSS, and JavaScript. MetagRoot also utilized R for statistical calculations and data analysis, all within the Model-View-Controller (MVC) pattern. The system is divided into Models, Views, Controllers, Services, and Factories. Models, represented by C# classes, define data and apply validation rules. Views use Razor (.cshtml) templates and C# logic to format data and generate HTML. Controllers act as intermediaries between Models and Views. Services manage core application logic and data operations, whereas Factories create specific components to minimize dependencies. When a user sends a request, the Controller identifies the required action, uses a Factory to generate the necessary Model, and retrieves data through Services. Once the Controller has organized the data, it passes everything to the View for HTML rendering, thereby completing the request-response cycle.

Finally, the application comes with a suite of integrated plugins and viewers. The integrated components include Diamond (38), which is utilized for rapid sequence alignment; the nextProt Feature Viewer (39), employed for the visualization of annotated sequence features; UpSet plots for showing intersections and unions, MSAViewer (40), used to display multiple sequence alignments; the SkyLign sequence logo viewer (41) for visualizing HMM models and frequencies, and the Mol* structure viewer (42), implemented for interactive 3D molecular structure visualization. Additionally, OpenStreetMap is incorporated to support geospatial data representation, and HMMER (43) is used for HMM-based sequence homology searches.

## Results

### Database components

MetaGroot is a specialized repository cataloging 71,091 protein families linked to plant root microbiomes, each comprising ≥100 sequences derived from integrated metagenomic (1,199 datasets), metatranscriptomic (327 datasets), and 2,978 reference genomes. These families analyzed across 29 million scaffolds and 421 million protein sequences, are enriched with annotations including multiple sequence alignments, Hidden Markov Models (HMMs), taxonomic/functional classifications, CRISPR elements, and ecosystem metadata (e.g., rhizosphere, nodules). Structural insights are provided via AlphaFold2-predicted 3D models, with 59,059 high-quality and 9,036 medium-quality structures.

### The metaGroot Interface

The metaGroot platform is accessed via a top-of-page navigation bar that provides main menus such as Browse, Sequence Search Tools, Statistics, and Downloads, ensuring access to every feature from any page (Figure 2).

**Figure 2.**
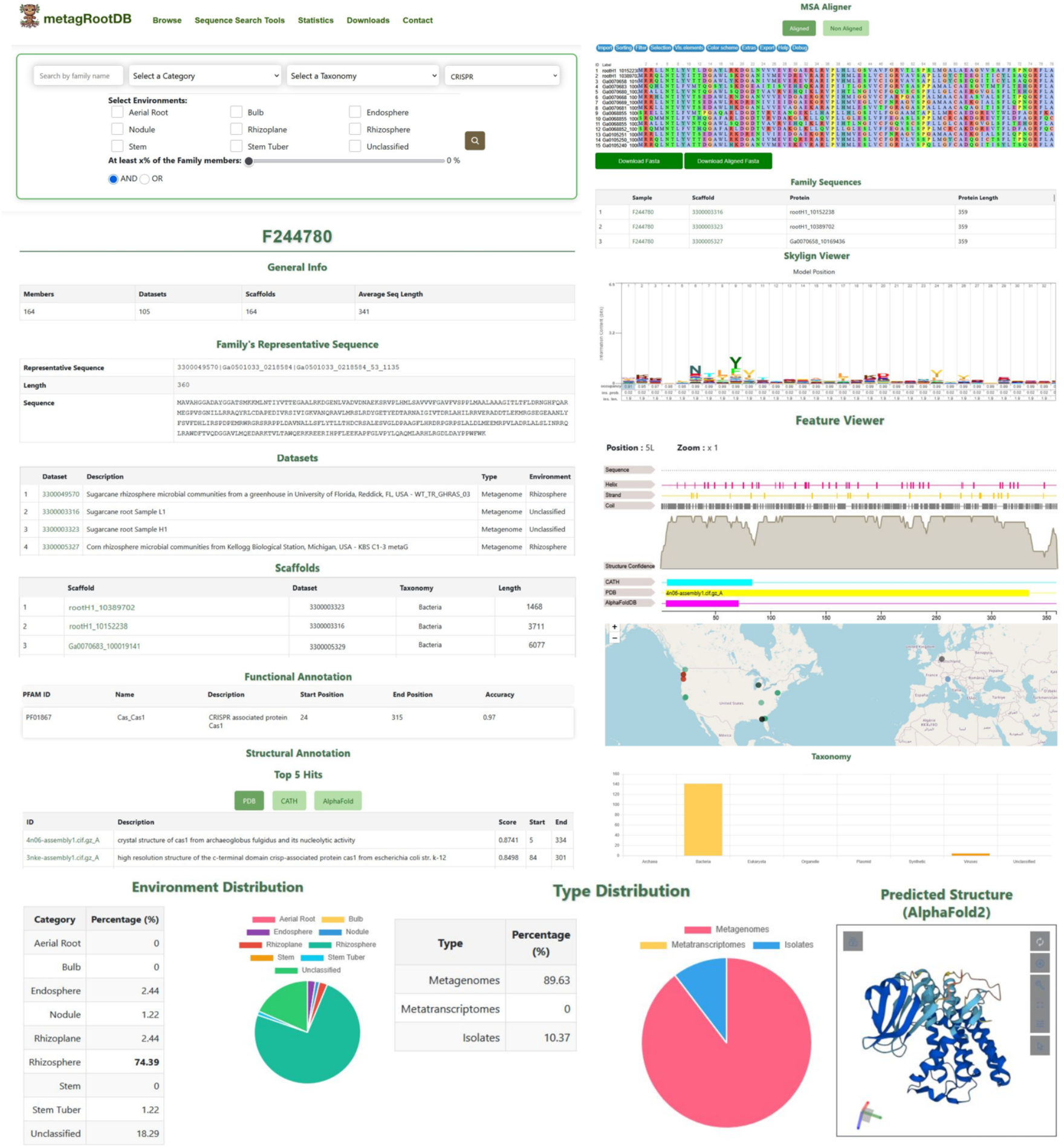
MetaGroot’s interface and summaries. The Browse Families page offers free-text search and filters for sequence category, taxonomy, ecosystem, and family size. Selecting a family displays its identifier, member, and dataset counts, average sequence length, and representative amino-acid sequence, followed by tables of associated Plant Datasets and scaffolds, Pfam functional domains, and top structural matches. Interactive viewers include a multiple sequence alignment with the customization and FASTA export, a sequence logo from the profile HMM with export options, a linear feature map of topology, secondary structure, conservation and domain annotations, and an embedded AlphaFold2 3D model. Metadata panels summarize phylogenetic assignments, gene-neighborhood contexts, ecosystem and data-type distributions, and global sampling locations. All alignments, profiles, models, and tables are downloadable in standard bioinformatics formats.

### Browse Families and Search Filters

Under Browse → Families, users encounter a comprehensive query interface that supports both simple and advanced searches. A free-text box accepts metaGroot family identifiers or keywords linked to family identifiers, while dropdown menus allow filtering by sequence category (Metagenome-only, Metatranscriptome-only, Mixed, or all families), taxonomy, and sequence type (e.g., CRISPR). Ecosystem checkboxes (Aerial Root, Bulb, Endosphere, Nodule, Rhizoplane, Rhizosphere, Stem, Stem Tuber, Unclassified) further narrow results, and a minimum-members slider is used for the retrieval of families with at least a percentage of their members coming from a specific habitat. The search options are further combined with an AND/OR logic toggle. Finally, users can constrain families by size before initiating the search. Once the criteria are set, a prominent search button retrieves matching families for online browsing or export.

### Family Entry: Data, Visualization, and Metadata Context

When a family is selected, the interface presents its unique identifier along with key summary statistics such as the total number of member sequences, contributing Plant Datasets, scaffolds, and the average sequence length. The representative sequence from the multiple alignment is displayed in full (complete with its amino-acid string and length) followed by comprehensive tables listing every associated Plant Dataset (including dataset ID, environmental metadata, and designation as metagenome or metatranscriptome) and every scaffold (showing scaffold ID, source dataset linkage, taxonomic assignment, and sequence length). All identifiers are linked to IMG/M.

Functional annotation is provided through Pfam domain hits, each annotated with start and end positions and per-domain confidence scores. Structural annotation reports the top five PDB, CATH, or AlphaFoldDB matches for the family’s representative sequence, listing TM-scores and alignment spans to convey fold similarity.

In addition, a suite of interactive tools enables in-depth exploration of these data. The multiple sequence alignment (MSA) can be viewed in “full” or “seed” mode, with options to apply custom coloring schemes, set identity and occupancy thresholds, search by motif or regular expression, and download the alignment in FASTA format. An adjacent sequence-logo viewer renders the family’s profile HMM as an interactive plot of residue probabilities (each position being clickable to reveal detailed scores). The underlying HMM can be exported in HMMER or HH-suite format. Per-residue predictions for transmembrane topology, secondary structure elements, conservation histograms, and Pfam (or other) domain annotations are mapped directly onto the representative sequence in a linear feature viewer. An AlphaFold2 model completes the picture when available by providing a rotatable, three-dimensional ribbon diagram of the predicted structure.

To contextualize the family ecologically and taxonomically, metaGroot integrates several metadata summaries. Phylogenetic assignments aggregated across all scaffolds reveal the family’s higher-level taxonomic composition, while a gene-neighborhood overview highlights conserved Pfam domains in the genomic vicinity of each family member. Ecosystem distributions are quantified in both a table and a pie chart, showing the percentage of sequences derived from each habitat category. A second pie chart breaks down members by data type (metagenome, metatranscriptome, isolate) and a world map plots the geographic coordinates of all sampling sites. Finally, a bar chart summarizes the family’s taxonomic breadth at the phylum or class level.

### Data Accessibility & Export

Every alignment, profile, annotation table, visualization, and structural model within metaGroot is fully exportable. Sequence alignments and family HMMs can be downloaded in standard FASTA and HMMER/HH-suite formats, respectively. The three-dimensional models are available in PDB format and metadata tables may be exported as CSV or TSV files.

### Sequence Search & Visualization Tools

Beyond browsing families, the Sequence Search Tools menu provides integrated HMMER and DIAMOND pipelines for querying the entire metaGroot database with user-supplied sequences or profile hidden Markov models. Users can refine search results by taxonomy or ecosystem, adjust sensitivity parameters such as cut-offs, substitution matrices, and gap costs, and then download tabulated hits in plain text format. A dedicated Pattern Search interface supports PROSITE-style motif queries or arbitrary regular expressions, enabling rapid detection of conserved sequence features across all family members.

In conclusion, metaGroot integrates data on plant root microbiome protein families by combining hundreds of metagenomic and metatranscriptomic datasets with thousands of curated reference genomes. Its user-friendly web platform enables researchers to explore 71,091 protein families through multiple criteria (e.g., sequence, structure, taxonomy, or ecosystem) while offering tools to visualize, and download alignments, HMM profiles, and AlphaFold2-predicted models. Comprehensive metadata, including CRISPR elements and geographic sampling data, further enriches analysis. In contrast, advanced search utilities (HMMER, DIAMOND, PROSITE/regex) with interactive visualization and customizable export features, accelerate hypothesis development around root-associated proteins. By bridging cutting-edge computational tools with actionable insights, metaGroot emerges as a pivotal resource for unraveling the molecular mechanisms of plant-microbe interactions and steering experimental and biotechnological advances in plant microbiome studies.

### Case Studies

#### Case 1: Integrating sequence and structural evidence for enhanced annotation precision

CRISPR elements are naturally occurring sequences in prokaryotic genomes that function as an adaptive immune system, recognizing and cleaving specific foreign genetic material through guide RNA and associated Cas enzymes. These elements have been repurposed as powerful tools in genetic engineering, enabling precise editing of DNA sequences to modify genes, correct mutations, or engineer organisms for research, medical, and industrial applications. Family F244780 represents a CRISPR-associated protein family, consisting of 164 protein sequences and 164 scaffolds, is shared across 5 habitats (*Endosphere, Nodule, Rhizoplane, Rhizosphere, Stem Tuber -*and *Unclassified*-), and is composed of approximately 10% reference genomes, with the remaining 90% sourced from metagenomic or metatranscriptomic datasets. Initial sequence analysis revealed a single Cas1 domain (Pfam: PF01867). In addition, structural searches provided enhanced functional insights. Key findings include a resolved Cas1 endonuclease structure (PDB: 4n06-assembly1) and a predictive CRISPR-associated endonuclease Cas1 model generated by AlphaFoldDB (AF-A0A645DPD0-F1-v4). CATH homology analysis further supported these findings by linking the domain to conserved structural folds indicative of its integration into a multi-component Cas1 protein complex (af_P9WPJ5_2_80_3.100.10.20). Structural hits of the F244780 family to PDB, CATH, and AlphaFoldDB highlight how structural insights can enhance or strengthen Pfam annotations, demonstrating that integrating structural and sequence data can improve predictive accuracy and resolve functional ambiguities. Indicative scaffolds for this family and structural models are shown in Figure 3A.

**Figure 3.**
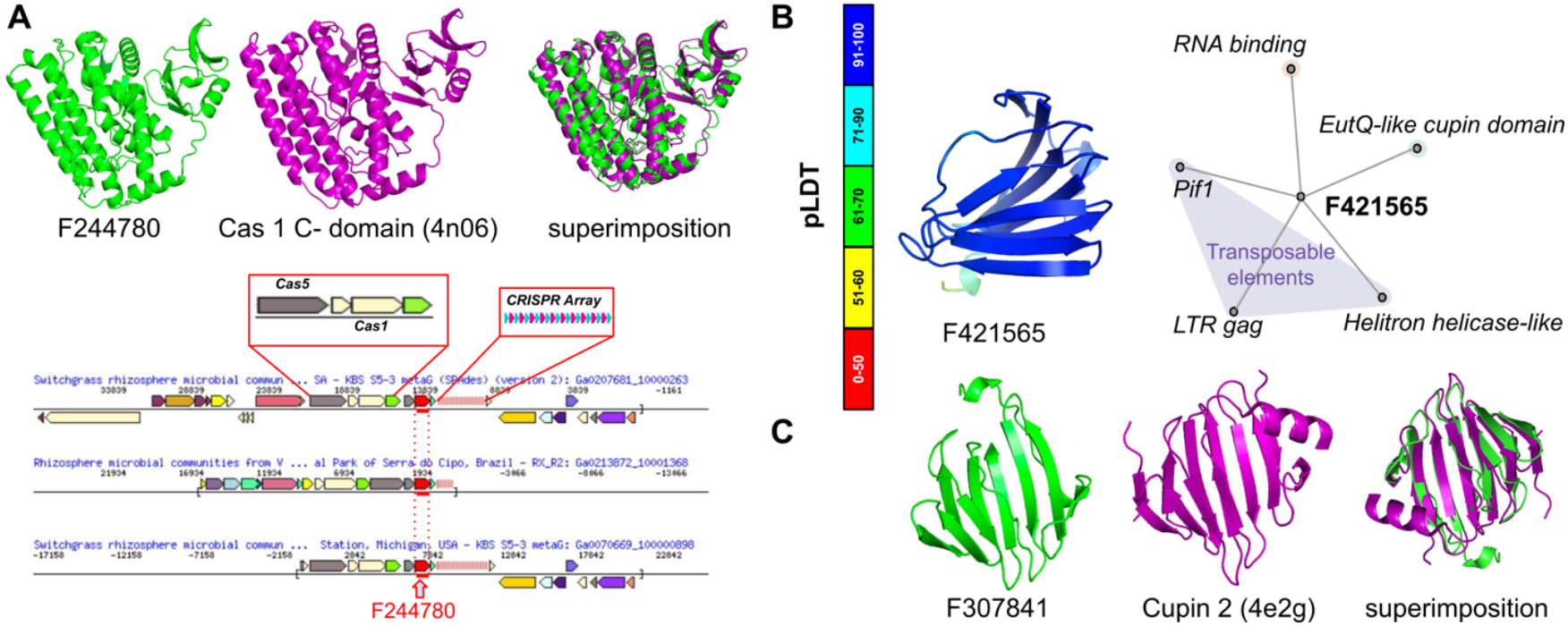
Example cases. **(A)** Family F244780 (green color) presents structural alignment with the CATH domain 4n06 (magenta color), achieving an alignment score of 0.9215. The superimposition highlights the structural similarity between this protein family and the CATH 4n06 domain. At the bottom, three scaffolds are displayed, emphasizing the presence of Cas domains and CRISPR arrays. **(B)** The family model F421565 shows no significant hits to CATH, PDB, or AlphaFoldDB, suggesting it may represent a potential novel fold, with pLDDT indicating the per-residue confidence score. On the left, a network generated by NORMA illustrates the gene neighborhood of the family, organized into annotated categories based on Gene Ontology and Pfam descriptions. The mobile genetic elements category (purple color) includes the Pif1, LTR gag, and Helitron helicase-like domain. Another category features proteins of RNA binding and EutQ-like cupin domain. **(C)** Family F307841 (green color) presents structural alignment with the CATH Cupin 2 domain (magenta color), achieving an alignment score of 0.8767. The superimposition highlights the structural similarity between this protein family and the CATH Cupin 2 (4e2g) domain.

#### Case 2: Identification of novel families

Novel protein families, defined by stringent sequence similarity thresholds (≤30% identity, ≤80% bidirectional query-subject coverage against reference proteomes or Pfam entries), represent a critical component of the “microbial dark matter.” These families are hypothesized to encode unexplored functional and evolutionary diversity, as they lack detectable sequence or structural homology to annotated proteins. To identify such families, those composed exclusively of metagenomic sequences with no Pfam matches were further analyzed against the NMPfamsDB catalog. For instance, family F421565, comprising 359 protein sequences and 307 scaffolds, is uniquely associated with the Nodule environment subtype (Figure 3B) and aligns with two NMPFamDB entries (F061583 and F098768). Looking at gene neighbors within the scaffold (visualized as a network by NORMA (44, 45)), the genomic context of this family reveals features characteristic of symbiosis-associated loci. Mobile element domains, including Helitron helicase-like, Pif1, and LTR gag, suggest a horizontal gene transfer origin. HGT (horizontal gene transfer) is a hallmark of symbiotic genomic islands observed in systems such as Bradyrhizobium japonicum, where HGT-derived loci mediate host-specific nitrogen fixation. The co-occurrence of RNA-binding domains (APO_RNA-binding, GO:”RNA_binding”) implies regulatory roles, potentially enabling post-transcriptional modulation under nodule-specific conditions. EutQ-like cupin domains, further point to metabolic adaptations tailored to the nodule niche. Together, the integration of mobile elements, regulatory components, and niche-specific metabolic functions supports the hypothesis that this locus facilitates nodule colonization or symbiotic interaction. These findings underscore the broader significance of microbial dark matter in elucidating uncharacterized mechanisms underlying host-microbe symbiosis, highlighting its potential to expand our understanding of functional innovation in uncultivated environmental niches.

#### Case 3: Functional annotation of uncharacterized reference genomes within a taxonomic family enriched through metagenomic and structural analysis

Integrating metagenomic data with reference genomes within the same taxonomic family is critical for comprehensive functional annotation, as metagenomes often provide indispensable insights into protein functions when reference genomes remain uncharacterized. This principle is exemplified through the analysis of family F307841, a phylogenetically cohesive group comprising 200 protein sequences and 196 scaffolds distributed across four distinct habitats (*Bulb, Endosphere, Rhizoplane, Rhizosphere -*and *Unclassified-*). The family comprises approximately 2.0% reference genomes, 3.5% metatranscriptomes, and 94.5% metagenomic sequences. Notably, no prior functional annotations in the Pfam database were associated with the reference genome sequences in this family. However, through metagenomic analysis, 22 sequences were identified as containing a conserved Cupin domain (PF07883), a structural motif frequently linked to metabolic or metal-binding functions. This annotation was validated by structural alignment to the crystal structure of Sthe2323 (PDB: 4e2g), a Cupin fold protein from Sphaerobacter thermophilus (Figure 3C).

## Discussion

The properties of the protein families identified in this study were initially examined for several patterns (Figure 4A). In the near-normal distribution of protein lengths, most proteins spanned 200 to 450 aminoacids, whereas a clear peak was observed between the range of 250 and 300. In addition, protein family sizes were skewed towards smaller groups, as many contained, on average, 150 to 300 members. In the case of metagenomes and metatranscriptomes, scaffold lengths were found to lie predominantly between the range of 1,000 and 5,000 base pairs. Lastly, the histogram illustrating the protein members per cluster underscored a tendency for these proteins to cluster mainly between 150 and 500 sequences. The biome distribution of protein families illustrated in the UpSet plot revealed significant variability across different plant-associated habitats (Figure 4B). One noteworthy observation was the prevalence of protein families found only in certain biomes (Rhizosphere and Nodule), suggesting unique functional adaptations. The rhizosphere accounted for the largest share of these exclusive families, aligning with its dynamic role in plant-microbe relationships. Likewise, a set of families (>25,000 families) is shared between the endosphere, rhizosphere, and rhizoplane, hinting at the specialized demands within internal tissues and along root surfaces, and reflecting shared functions tied to root interactions and resource uptake. In contrast, habitats like bulbs and nodules demonstrated fewer overlaps, maybe due to their unique physiology and characteristics. Bulbs, unlike the other environments discussed, are storage organs characterized by layers in their structure, high amounts of sugars, and low oxygen. Similarly, nodules are formed due to symbiotic interaction between the plant and nitrogen-fixing bacteria, and as a result, they provide an environment favorable for nitrogenase (46).

**Figure 4.**
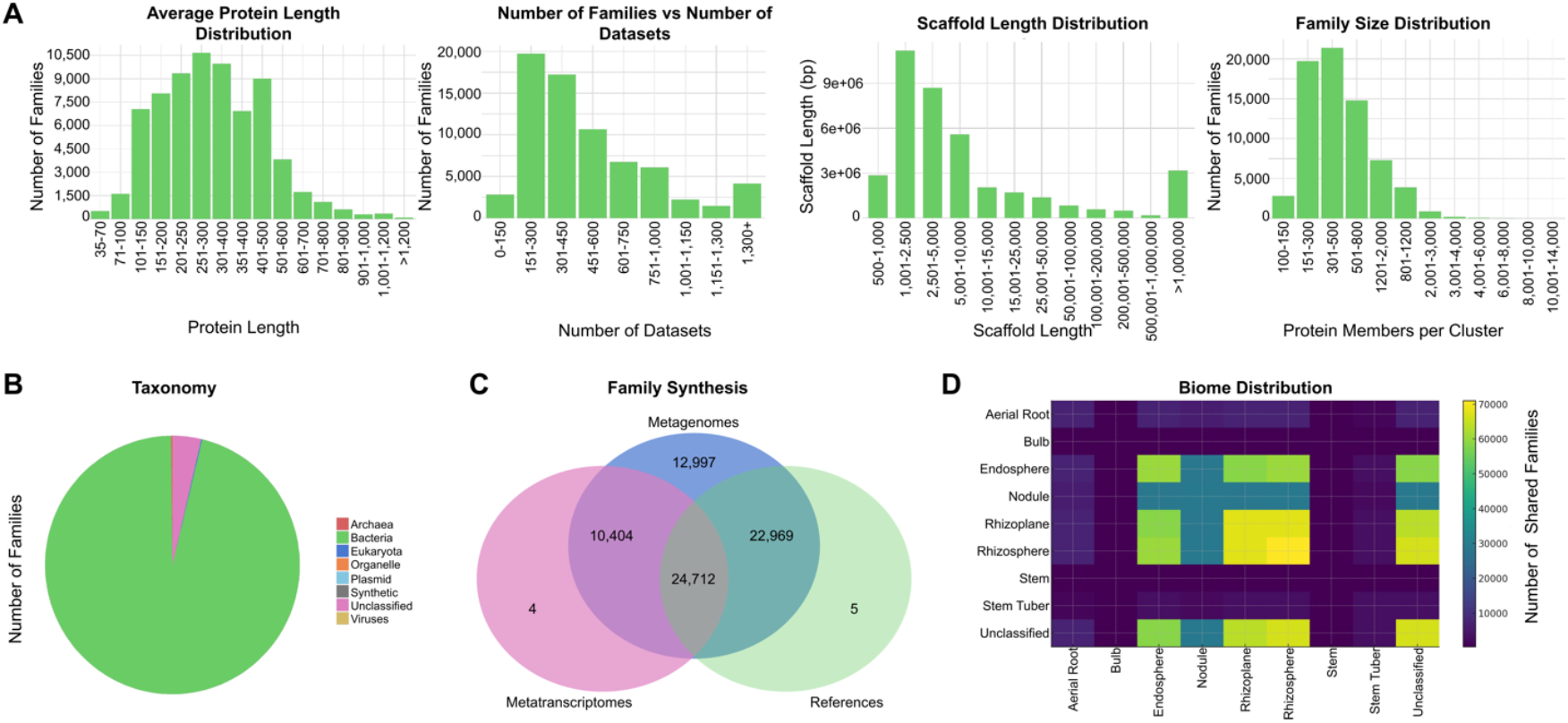
Distribution and characteristics of protein families, biomes, taxonomy, and habitats. **(A)** Histograms displaying average protein lengths, number of datasets, scaffold length distributions, and protein members. **(B)** Taxonomic composition of sequences, with bacterial taxa representing the overwhelming majority, followed by unclassified families, a few archaeal, and minimal eukaryotic and viral sequences. **(C)** Venn diagram demonstrating overlap among metagenomic, metatranscriptomic, and reference genome datasets. **(D)** Heatmap showing the pairwise family overlap across the various habitats.

In terms of taxonomy (Figure 4C), bacterial taxa were determined to constitute the vast majority (approximately 94,91%) of all classified sequences. Archaeal sequences, fewer in number, accounted for a minority (0.21%). Conversely, eukaryotic and viral taxa, as well as organelles, plasmids, and synthetic categories, represented only a small fraction of the dataset (0.21%), whereas 4.67% remained unclassified. This taxonomic distribution underscores the bacterial-centric nature of the microbial communities represented while emphasizing that archaea, though present, comprise only a small proportion relative to bacteria.

Lastly, the overlap among metagenomes, metatranscriptomes, and reference genomes is shown in a Venn diagram (Figure 4D). A substantial core consisting of 24,712 sequences was shared across all three categories. Pairwise intersections were also significant, notably between metagenomes and reference genomes (22,969 sequences), as well as between metagenomes and metatranscriptomes (10,404 sequences). Metatranscriptomes and reference genomes, however, exhibited minimal exclusivity, containing only four and five unique sequences, respectively. Thus, metagenomic datasets effectively encompass most sequence diversity, capturing sequences found within reference genomes and metatranscriptomic datasets. Collectively, these results depict an ecosystem dominated by bacterial sequences characterized by predominantly small protein clusters across certain habitats.

In conclusion, metagRoot represents a significant advancement in the study of plant root microbiomes by integrating metagenomic, metatranscriptomic, and reference genome data into a unified, richly annotated resource. By cataloging 71,091 protein families with structural, functional, and ecological context, metagRoot bridges a critical gap in existing databases, enabling researchers to explore the molecular mechanisms underpinning plant-microbe interactions with unprecedented depth. The inclusion of AlphaFold2-predicted 3D models, CRISPR elements, habitat-specific metadata, and interactive analytical tools empowers users to uncover novel protein functions, refine annotations, and investigate evolutionary and ecological dynamics across root-associated niches. As a publicly accessible platform, metagRoot not only accelerates hypothesis-driven research but also fosters microbiome-informed innovations in agriculture.

## Data Availability

All data, including sequences, multiple sequence alignments, 3D models, and the associated metadata, can be found at: https://doi.org/10.5281/zenodo.15488157.

